# An ACE2-dependent Sarbecovirus in Russian bats is resistant to SARS-CoV-2 vaccines

**DOI:** 10.1101/2021.12.05.471310

**Authors:** Stephanie N. Seifert, Shuangyi Bai, Stephen Fawcett, Elizabeth B. Norton, Kevin J. Zwezdaryk, James Robinson, Bronwyn Gunn, Michael C. Letko

## Abstract

Spillover of sarbecoviruses from animals to humans has resulted in outbreaks of severe acute respiratory syndrome SARS-CoVs and the ongoing COVID-19 pandemic. Efforts to identify the origins of SARS-CoV-1 and −2 has resulted in the discovery of numerous animal sarbecoviruses – the majority of which are only distantly related to known human pathogens and do not infect human cells. The receptor binding domain (RBD) on sarbecoviruses engages receptor molecules on the host cell and mediates cell invasion. Here, we tested the receptor tropism and serological cross reactivity for RBDs from two sarbecoviruses found in Russian horseshoe bats. While these two viruses are in a viral lineage distinct from SARS-CoV-1 and −2, one virus, Khosta-2, was capable of using human ACE2 to facilitate cell entry. Viral pseudotypes with a recombinant, SARS-CoV-2 spike encoding for the Khosta 2 RBD were resistant to both SARS-CoV-2 monoclonal antibodies and serum from individuals vaccinated for SARS-CoV-2. Our findings further demonstrate that sarbecoviruses circulating in wildlife outside of Asia also pose a threat to global health and ongoing vaccine campaigns against SARS-CoV-2

**ONE SENTENCE SUMMARY:** European bat coronaviruses that are only distantly related to SARS-CoV-2 but use the same cell entry route, escape the immune response against SARS-CoV-2 vaccines, driving the need for broader vaccines.

## Introduction

Zoonotic spillover of sarbecoviruses from animals to humans has led to the emergence of highly pathogenic human viruses, SARS-CoV-1 and −2, with the later leading to the largest global pandemic in modern history. Researchers around the world are ramping up the pace of viral discovery efforts, expanding the sequence databases with new animal sarbecoviruses in circulation. While some laboratory experiments have been performed with these new viruses, demonstrating a range of host tropisms, several viruses remain untested, and thus their ability to transmit to humans is unknown.

Coronaviruses are covered with a spike glycoprotein (S) that engages with receptor molecules on the surface of host cells and mediates viral infection of the cell. A small region within the spike proteins of sarbecoviruses, known as the receptor binding domain (RBD), contains all of the structural information necessary to engage with the host receptor. We and others have experimentally classified the majority of published sarbecovirus RBDs into different clades based on sequence and functional data: clade 1, identified in Asian bats, contains no deletions and binds to host receptor, Angiotensin-Converting Enzyme 2 (ACE2), whereas clade 2, also identified in Asian bats, contains 2 deletions and does not use ACE2 and clade 3 viruses, found more widely in Africa and Europe, contain 1 deletion and some can infect using an unknown receptor, while other clade 3 RBDs use ACE2(*1*–*3*). Recently, several viruses were identified in China that comprise a fourth clade that also interact with human ACE2(*4*).

In late 2020, two clade 3 sarbecoviruses were identified in Rhinolophus bats in Russia: Khosta-1 was found in *Rhinolophus ferrumequinum* and Khosta-2 in *R. hipposideros(5)*. Similar to other European and African clade 3 viruses, the Khosta viruses are divergent from the RBD found in SARS-CoV-1 and −2. Recently, the Khosta 2 RBD was shown in biochemical assays to bind human ACE2 (*6*). Here, we used pseudotype entry assays to assess the receptor preference of the RBDs as well as the full-length spikes from the Khosta viruses and confirm that Khosta 2 spike can readily use human ACE2 to infect cells. We also assessed the antibody neutralization of a chimeric SARS-CoV-2 spike encoding for the Khosta2 RBD to assess the protection offered by current SARS-CoV-2 based vaccines against future emerging Sarbecovirus threats. Our findings highlight the urgent need to continue development of new, and broader-protecting Sarbecovirus vaccines.

## Results

### Khosta virus receptor binding domains are distinct from human viruses

Khosta-1 and −2 were identified by Alkhovsky and colleagues in bat samples collected between March-October 2020 near Sochi National Park(*5*). Phylogenetic analysis of the conserved viral gene, Orf1ab, revealed these viruses were most closely related to another sarbecovirus found in Bulgaria in 2008 (known as BM48-31 or Bg08), and form a lineage sarbecoviruses distinct from human pathogens, SARS-CoV-1 and - 2(*5*). A list of viruses and accession numbers used in this study can be found in table 1. Phylogenetic analysis of the spike RBD further reflected the close relatedness between Khosta −1 and −2 with BM48-31 and other clade 3 RBD viruses we have previously tested from Uganda and Rwanda(*1*, *7*) (Fig. 1A). Clade 3 RBDs, including the Khosta viruses, contain a truncated surface-exposed loop, as compared to the ACE2-dependent, clade 1 viruses such as SARS-CoV and additionally vary in most of the residues known for clade 1 viruses to interact with ACE2(*1*, *2*, *7*, *8*).

**Figure 1.**
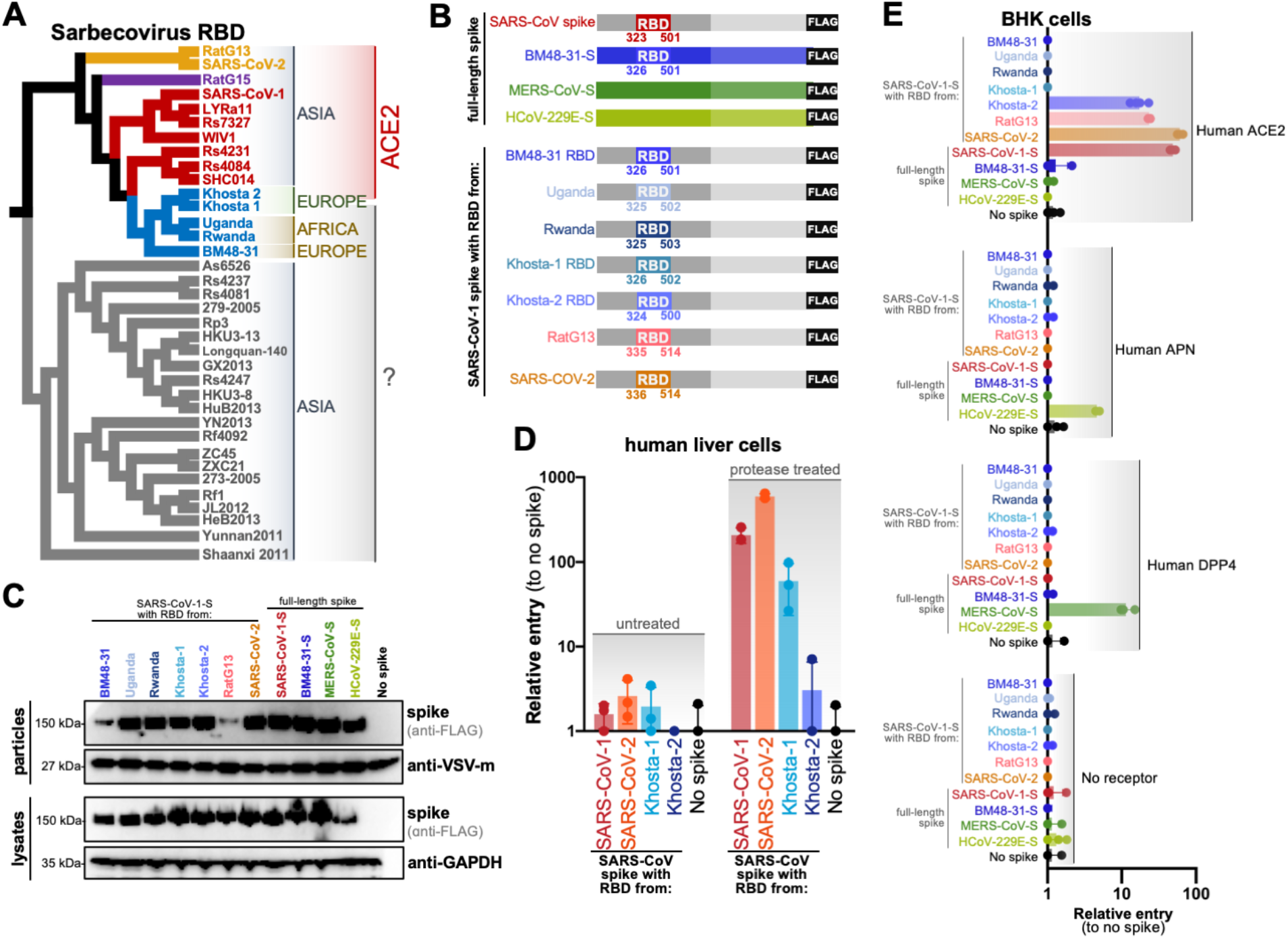
Khosta-2 uses human ACE2 to infect cells. (**A**) Sarbecovirus Receptor Binding Domain Cladogram based on amino acid sequenced. Countries of origin and known host receptors are indicated to the right. Clade 1 viruses are shown in red and orange, clade 2 in grey, clade 3 in blue and clade 4 in purple. (**B**) Diagram of spike constructs used for this study. The SARS-CoV-1 RBD was replaced with RBDs from other sarbecoviruses. (**C**) Expression and incorporation of viral pseudotypes by westernblot. (**D**) Huh-7 cells were infected with pseudotypes in the presence of absence of trypsin. Cells were infected in triplicate. (**E**) BHK cells were transfected with receptors and infected in the absence of trypsin. Cells were infected in quadruplicate.

**Table 1.** Sarbecovirus sequences used in this study.

|  | **Virus** | **Accession** | **Clade** | **Host species** | **Location** |
| --- | --- | --- | --- | --- | --- |
| 1 | SARS Urbani | AY278741 | 1 | human | Guangdong, China (origin of outbreak) |
| 2 | WIV1 | KF367457 | 1 | *Rhinolophus sinicus* | Yunnan, China |
| 3 | LYRa11 | KF569996 | 1 | *Rhinolophus affinis* | Baoshan, Yunnan, China |
| 4 | Rs7327 | KY417151 | 1 | *Rhinolophus sinicus* | Kunming, Yunnan Province, China |
| 5 | Rs4231 | KY417146 | 1 | *Rhinolophus sinicus* | Kunming, Yunnan Province, China |
| 6 | Rs4084 | KY417144 | 1 | *Rhinolophus sinicus* | Kunming, Yunnan Province, China |
| 7 | RsSHC014 | KC881005 | 1 | *Rhinolophus sinicus* | Yunnan, China |
| 8 | SARS-CoV-2 | MN908947 | 1 | human | Wuhan, Hubei, China |
| 9 | RatG13 | MN996532 | 1 | *Rhinolophus affinis* | Yunnan, China |
| 10 | RatG15 | from preprint | 4 | *Rhinolophus affinis* | Mojiang County, Yunnan Province, China |
| 11 | As6526 | KY417142 | 2 | *Aselliscus stoliczkanus* | Kunming, Yunnan Province, China |
| 12 | Yunnan2011 | JX993988 | 2 | *Chaerephon plicata* | Yunnan, China |
| 13 | Shaanxi2011 | JX993987 | 2 | *Rhinolophus pusillus* | Shaanxi, China |
| 14 | 279-2005 | DQ648857 | 2 | *Rhinolophus macrotis* | Hubei, China |
| 15 | Rs4237 | KY417147 | 2 | *Rhinolophus sinicus* | Kunming, Yunnan Province, China |
| 16 | Rs4081 | KY417143 | 2 | *Rhinolophus sinicus* | Kunming, Yunnan Province, China |
| 17 | Rp3 | DQ071615 | 2 | *Rhinolophus pearsoni* | Nanning, Guangxi, China |
| 18 | Rs4247 | KY417148 | 2 | *Rhinolophus sinicus* | Kunming, Yunnan Province, China |
| 19 | HKU3-8 | GQ153543 | 2 | *Rhinolophus sinicus* | Guangdong, China |
| 20 | HKU3-13 | GQ153548 | 2 | *Rhinolophus sinicus* | Guangdong, China |
| 21 | GX2013 | KJ473815 | 2 | *Rhinolophus sinicus* | Guangxi, China |
| 22 | Longquan-140 | KF294457 | 2 | *Rhinolophus monoceros* | Longquan, Zhejiang, China |
| 23 | YN2013 | KJ473816 | 2 | *Rhinolophus sinicus* | Yunnan, China |
| 24 | Rf4092 | KY417145 | 2 | *Rhinolophus ferrumequinum* | Kunming, Yunnan Province, China |
| 25 | ZXC21 | MG772934 | 2 | *Rhinolophus sinicus* | Zhoushan City, Zhejiang, China |
| 26 | ZC45 | MG772933 | 2 | *Rhinolophus sinicus* | Zhoushan City, Zhejiang, China |
| 27 | JL2012 | KJ473811 | 2 | *Rhinolophus ferrumequinum* | Jilin, China |
| 28 | HuB2013 | KJ473814 | 2 | *Rhinolophus sinicus* | Hubei, China |
| 29 | Rf1 | DQ412042 | 2 | *Rhinolophus ferrumequinum* | Yichang, Hubei, China |
| 30 | HeB2013 | KJ473812 | 2 | *Rhinolophus ferrumequinum* | Hebei, China |
| 31 | 273-2005 | DQ648856 | 2 | *Rhinolophus ferrumequinum* | Hubei, China |
| 32 | BM48-31 | NC014470 | 3 | *Rhinolophus blasii* | Strandja Nature Park, Bulgaria |
| 33 | Uganda | MT726044 | 3 | *Rhinolophus ferrumequinum* | Uganda |
| 34 | Rwanda | MT726045 | 3 | *Rhinolophus ferrumequinum* | Rwanda |
| 35 | Khosta 1 | MZ190137 | 3 | *Rhinolophus ferrumequinum* | Greater Caucasus , Russia |
| 36 | Khosta 2 | MZ190138 | 3 | *Rhinolophus hipposideros* | Greater Caucasus , Russia |

### RBD from Khosta viruses mediate entry into human cells

Using our scalable Sarbecovirus RBD entry platform, we replaced the RBD from SARS-CoV-1 spike with the Khosta RBDs and generated chimeric spike expression plasmids (Fig. 1B)(*1*). For comparison, we also included chimeric RBD spikes for other clade 3 RBDs we have previously tested (BM48-31, Uganda, Rwanda) as well as SARS-CoV-2 and related RatG13 viruses. These chimeric spike expression constructs were used to produce BSL2-compatible viral reporter pseudotypes with Vesicular Stomatitis Virus expressing a dual GFP-luciferase reporter(*1*). All of the chimeric spike proteins expressed to similar levels in mammalian cells and incorporated in Vesicular Stomatitis Virus (VSV). Chimeric spike with the RBD from BM48-31 and RatG13 showed reduced incorporation but this did not correlate with viral entry phenotypes observed in later experiments (Fig. 1C, D, E).

To test human cell compatibility, we first infected the human liver cell line, Huh-7, with pseudotypes bearing the chimeric Khosta RBD spikes. In the absence of the addition of an exogenous protease, trypsin, the pseudotypes exhibited almost no entry in these cells, which has been observed for other sarbecoviruses and is attributed to low endogenous expression of ACE2. However, when trypsin was included during the infection, entry signal strongly increased for SARS-CoV-1 and −2 RBDs as well as the Khosta RBDs (Fig. 1D). As we and others have shown previously, trypsin enhancement of sarbecovirus entry is still receptor dependent, suggesting that the Khosta virus RBDs were using a receptor present in human cells to mediate infection(*1*, *9*).

### The RBD from Khosta-2 infects cells using human ACE2

To characterize potential receptors for the Khosta viruses, we performed a classic receptor tropism test, where we transfected Baby Hamster Kidney (BHK) cells with human orthologues of known coronavirus receptors and then infected with our pseudotype panel. Unlike 293T cells, which express low levels of human ACE2 and potentially other coronavirus receptors and have been shown to have low but measurable permissivity to SARS-CoV infection, BHK cells are generally considered completely non-permissive for sarbecoviruses unless a suitable receptor is supplemented(*10*). The Khosta-1 RBD failed to infect cells expressing any of the receptors, while Khosta 2 RBD clearly infected cells expressing human ACE2 (figure 1E). The level of cell entry mediated by the Khosta 2 RBD was similar to RatG13, a bat sarbecovirus with high similarity to SARS-CoV-2 in the RBD that also binds human ACE2, albeit with lower efficiency than the human pathogen(*11*, *12*) (Fig. 1E). In contrast to the ACE2 results, only the human virus, HCoV-229E, could infect cells expressing Aminopeptidase N (APN), and MERS-CoV spike could only infect cells expressing dipeptidyl peptidase IV (DPP4) – the known receptors for these viruses, demonstrating these receptors were expressed to functional levels (Fig. 1e).

### Full-length Khosta spikes infect human cells through ACE2

While the RBD from Khosta 2 can use human ACE2 in functional assays (Fig. 1) and bind ACE2 as a purified protein fragment(*6*), other domains in spike vary between the Khosta and SARS-CoV spikes. We had the full-length Khosta spike genes synthesized, generated viral pseudotypes and tested their infectivity on human cells (Fig. 2). Similar to the chimeric SARS-CoV-based spikes, full-legnth Khsota spikes could also infect Huh-7 cells in the presence of trypsin (Fig. 2A) and the Khosta 2 spike was capable of infecting 293T cells expressing human ACE2 even in the absence of trypsin (Fig. 2B). Analagous to what we have shown with other full-length Sarbecovirus spikes, the full-length Khosta 2 spike was less infectious than the chimeric SARS-CoV-based spike (Fig. 2B)(*1*).

**Figure 2.**
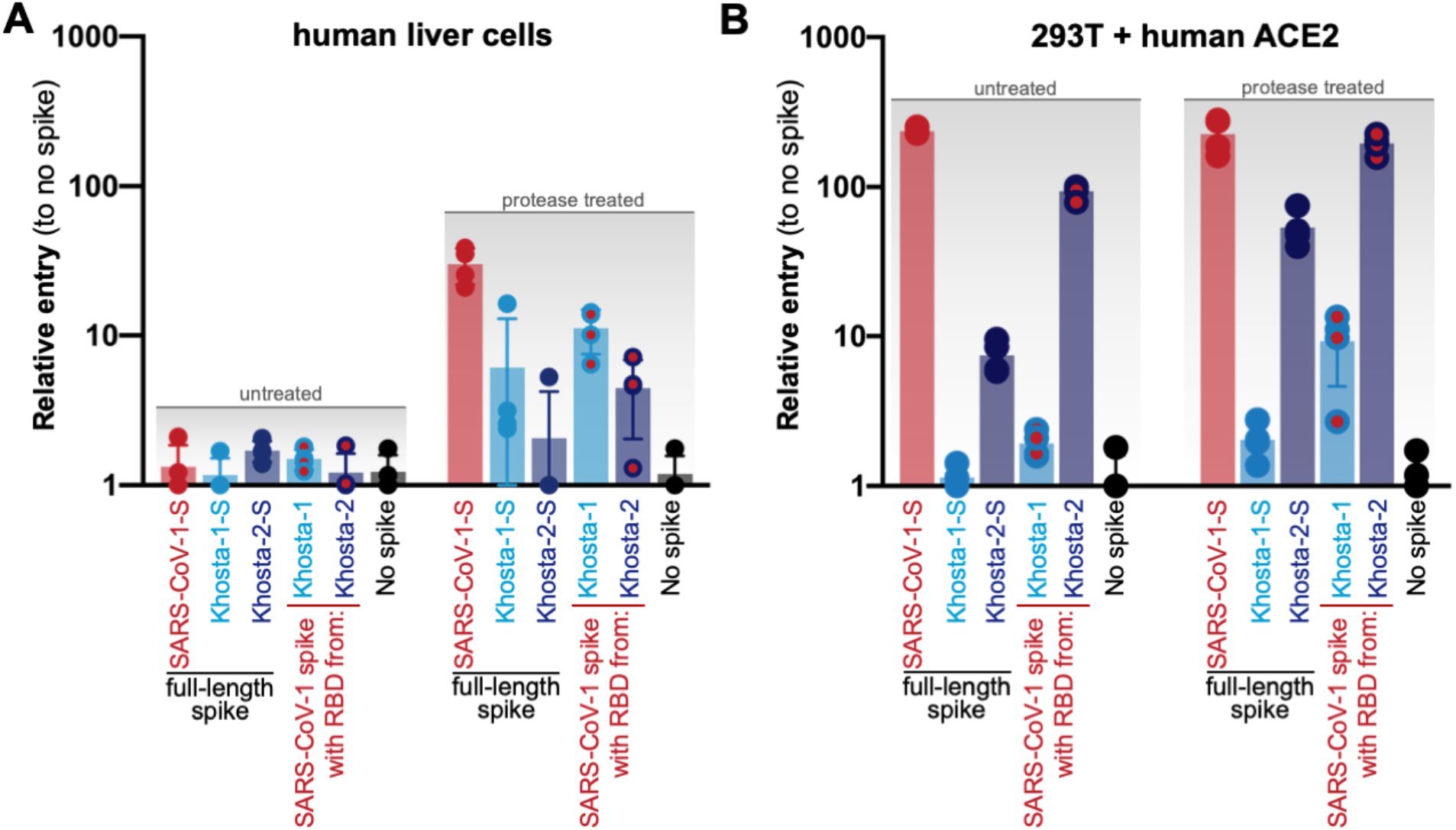
Full-length Khosta spikes infect human cells. (**A**) Huh-7 or (**B**) 293T cells that stably express human ACE2 cells were infected with VSV pseudotypes bearing the indicated spike protein.

### A SARS-CoV-2 based, Khosta2-chimeric spike is resistant to serum from SARS-CoV-2 vaccinated individuals

SARS-CoV-2 can infect a wide range of species and has now spilled back into both wild and domestic animals (*13*–*22*). Many animal species carry their own coronaviruses, and with the discovery of additional ACE2-dependent sarbecoviruses in broader geographic regions, the risk of new recombinant viruses is rising. To mimic the potential recombinant threat from the Khosta viruses, we generated VSV pseudotyped particles carrying a chimeric SARS-CoV-2-based spike with the RBD from the Khosta viruses (Fig. 3A). Similar to our earlier SARS-CoV-based spikes, the SARS-CoV-2 chimeric spikes were also infectious in 293T cells expressing human ACE2 (Fig. 3B). To assess if the ACE2-dependent Khosta 2 RBD and SARS-CoV-2 RBD were cross-reactive, we incubated pseudotyped particles with increasing amounts of the SARS-CoV-2 RBD-specific monoclonal antibody, Bamlanivimab. Surprisingly, while SARS-CoV-2 spike was effectively neutralized by the antibody, the SARS-CoV-2 spike with the Khosta 2 RBD was completely resistant, suggesting little cross-reactivity between the two RBDs (Fig. 3C). We repeated the pseudotype experiment using serum from vaccinated individuals and saw a similar trend: the wild-type SARS-CoV-2 spike was easily inhibited by serum from individuals who received either the Moderna or Pfizer vaccine, but the SARS-CoV-2-Khosta-2 RBD spike was resistant (Fig. 3D). At higher dilutions of serum, there was a reduction in the chimeric spike infectivity, but this was significantly less than the wildtype spike at similar serum concentrations (Fig. 3E). The Khosta 2 RBD shares approximately 60% similarity with various SARS-CoV-2 spikes at the amino acid level, which may explain their low cross-reactivity (Fig. 3F). Taken together, these results demonstrate that new recombinant sarbecoviruses may pose a threat to current SARS-CoV-2 vaccines.

**Figure 3.**
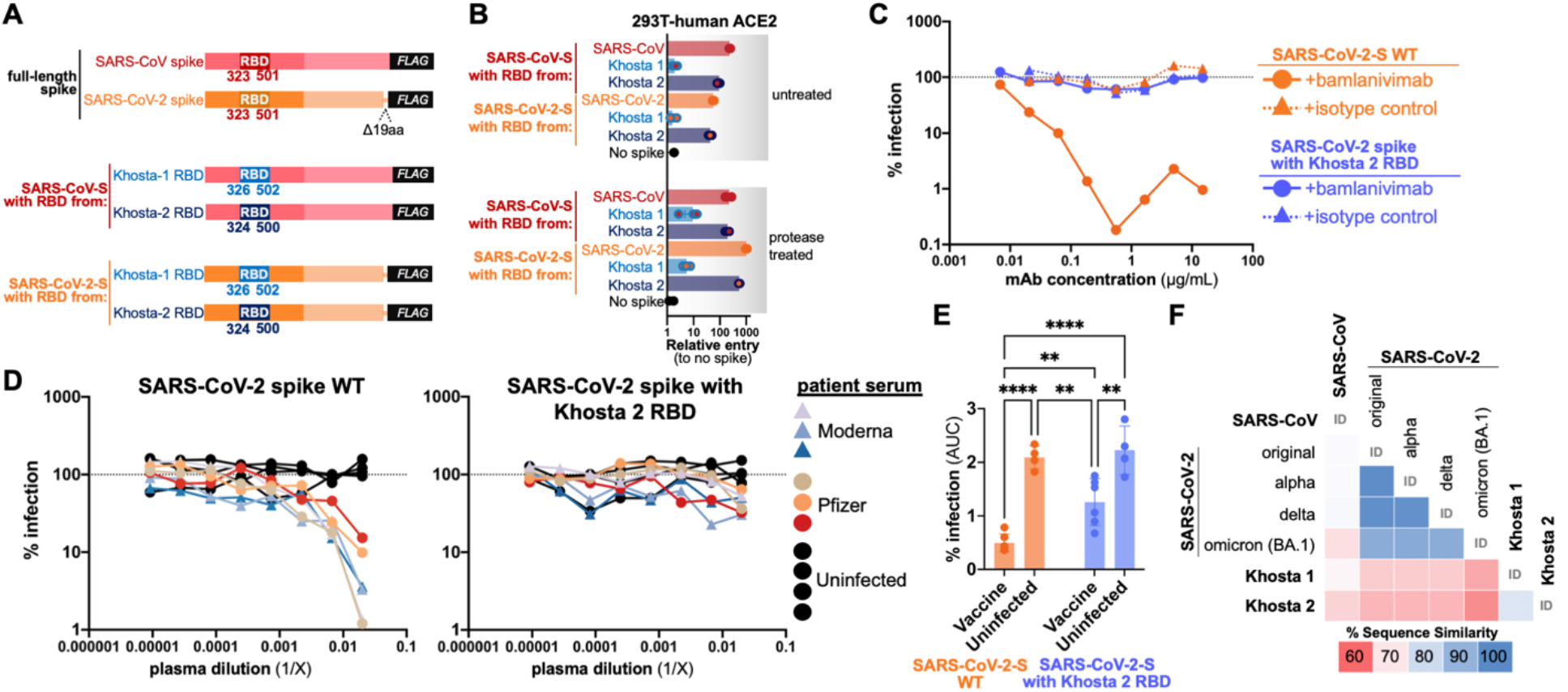
Chimeric SARS-CoV-2-Khosta 2 spike is resistant to current vaccines. (**A**) The RBD from SARS-Cov-2 spike was replaced with Khosta 2 RBD. (**B**) Pseudotpyes with indicated chimeric spikes were used to infect 293T cells stably expressing human ACE2. Pseudotypes were combined with (**C**) bamlanivimab or (**D**) vaccinated patient serum at various concentrations and used to infect 293T-hACE2 cells.(**E**) Area under the curve analysis for data in panel D. p-value 0.0021 (**), 0.0002 (***), <0.0001 (****) (**F**) Sequence identity matrix between Khosta 2 and known SARS-CoV-2 variants of concern

## Discussion

Khosta 1 and 2 viruses are most closely related to other clade 3 RBD viruses, which have been found across a much wider geographic range than the clade 1 viruses(*1*, *7*, *23*, *24*). As the researchers who initially discovered the Khosta viruses note with their findings: the Khosta bat sarbecoviruses are genetically distinct from human SARS-CoVs in that they lack genetic information encoding for some of the genes thought to antagonize the immune system and contribute to pathogenicity, such as Orf8(*5*). Unfortunately, because coronaviruses are known to recombine in co-infected hosts, the recent identification of SARS-CoV-2 spillover from humans back in wildlife populations opens the possibility of new human-compatible sarbecoviruses (*13*–*15*, *20*–*22*).

In the presence of trypsin, both Khosta-1 and −2 RBDs and spike were capable of infecting human cells, with Khosta-1 performing notably stronger than Khosta-2, however in our receptor-specific assay, only Khosta-2 could infect cells expressing human ACE2 without exogenous protease (Fig. 1D, 1E, Fig. 2). We have previously shown that a small number of clade 2 RBDs, such as As6526, also exhibit protease-mediated, ACE2-independent entry, and similar phenotypes have been described for other bat coronaviruses(*1*, *25*). Because not all of the clade 2 and 3 viruses exhibit this phenotype, these findings collectively suggest that some coronaviruses can infect human cells through a presently unknown receptor. Sarbecoviruses have been shown to co-circulate in bats, so this variation in receptor usage among closely related viruses may even represent an evolutionary strategy for viral persistence within the reservoir host population (*2*).

Current universal sarbecovirus vaccines in development include mostly clade 1 viruses and one of the clade 2 viruses but do not include any members from clade 3(*26*, *27*). Our results here suggest there is little cross-reactivity between clade 1 and clade 3 RBDs that use human ACE2 (Fig. 3C). More concerning was our observation that serum from vaccinated individuals was significantly less effective at neutralizing pseudotypes when just the SARS-CoV-2 RBD was replaced with the Khosta 2 RBD (Fig. 3D-E). These findings are not too surprising given that the Khosta 2 RBD only shares about 60% sequence identity with SARS-CoV-2, and the serum we tested was from individuals vaccinated only with RBD-specific vaccines from Moderna or Pfizer (Fig. 3E-F). Curiously, the Khosta 2 RBD is least similar to the currently circulating Omicron variant of SARS-CoV-2; with each new variant of concern decreasing in similarity to Khosta 2 (Fig. 3F). Given that natural infection or vaccination with a whole spike raises antibodies directed at other regions of spike, it is still possible that new sarbecoviruses or recombinant SARS-CoV-2 would be neutralized by serum from some individuals.

Our findings with chimeric, SARS-CoV-2 spike show that just replacing the RBD is sufficient to reduce efficacy of SARS-CoV-2 spike-directed vaccines (Fig. 3). However, sarbecovirus recombination in nature typically occurs via template switching resulting in acquisition of regions larger than the NTD (*28*). Thus, a naturally recombinant virus with Khosta 2 may actually acquire more Khosta 2 spike, which as we show here with full protein, is also infectious against human cells and ACE2 (Fig. 2). Taken together, our findings with the Khosta viruses underscore the urgent need to develop broader-protecting universal Sarbecovirus vaccines.

## Methods

### Phylogenetic analysis

Genbank accession numbers for all sarbecovirus spike sequences used in this study are available in table 1. Amino acid sequences for the receptor binding domain of the spike glycoprotein were aligned using ClustalW multiple sequence alignment with default parameters. A maximum likelihood phylogenetic tree was inferred with PhyML v. 3.0(*29*) using the ‘WAG’ matrix +G model of amino acid substitution as selected by Smart Model Selection method with 1000 bootstrap replicates(*30*). The final tree was then visualized as a cladogram with FigTree v1.4.4 (https://github.com/rambaut/figtree).

### Plasmids and sequences

Untagged human orthologues of ACE2 (Q9BYF1.2), APN (NP_001141.2), and DPP4 (XM_005246371.3) were described previously(*1*). Spike sequences from HCoV-229E (AB691763.1), MERS-CoV (JX869059.2), and SARS-CoV-1 (AY278741) were codon-optimized, appended with a carboxy-terminal FLAG tag sequence separated by a flexible poly-glycine linker and cloned into pcDNA3.1+ as previously described(*1*). SARS-CoV-2 spike (MN997409.1) was codon optimized, modified to including silent cloning sites flanking the RBD, and C-terminal 19 amino acid truncation was introduced to enhance pseudotyping (*1*, *31*). The SARS-CoV-1 RBD was removed with KpnI and XhoI, and the SARS-CoV-2 RBD was removed with BamHI and PflMI. Codon-optimized, synthesized RBD fragments were cloned into the spike backbones as previously described(*1*).

### Cells and pseudotype assay

293T, Huh-7 (human liver cells), and BHKs were maintained under standard cell culture conditions in DMEM with L-glutamine, antibiotics, and 10% Fetal Bovine Serum. Single-cycle, Vesicular Stomatitis Virus (VSV) pseudotype assays were performed as previously described(*1*). Briefly, 293T “producer cells” were transfected with spike plasmids or empty vector as a “no spike” control and infected 24-hours later with VSV-ΔG-luc/GFP particles, generating chimeric-spike pseudotyped particles that were harvested 72 hours post-transfection and stored at −80°C. Target cells were plated in 96-well format, and spin-infected in quadruplicate with equivalent volumes of viral pseudotypes at 1200xG for 1 hour at 4°C. Infected cells were incubated for approximately 18-20 hours and luciferase was measured using the Promega BrightGlo luciferase kit following manufacturers’ instructions (Promega). Entry signal was normalized to the average signal for the “no spike” control. Plates were measured and analyzed in triplicate. Data are representative of four complete biological replicates.

### Western blot

Viral pseudotypes were concentrated and 293T producer cells were lysed in 1% SDS and clarified as described previously(*1*). Lysates were analyzed on 10% Bis-Tris PAGE-gel (Thermo Fisher Scientific) and probed for FLAG (Sigma-Aldrich; A8592; 1:10000); GAPDH (Sigma-Aldrich, G8795, 1:10000); and/or VSV-M (Kerafast, 23H12, 1:5000). Signal was detected using SuperSignal West substrate (Thermo-Fisher).

### Patient serum samples

Deidentified plasma samples were from subjects recruited from the Greater New Orleans community under Tulane Biomedical Institutional Review Board (federalwide assurance number FWA00002055, under study number 2020-585).

### Monoclonal and serum neutralization assays

Sera from six vaccinated (3 Pfizer and 3 Moderna) and 4 infected donors were kindly provided by Tulane University. 293T-ACE2 cells were seeded at 2.5 x 10^4^ cells/well in 96-well format and grown for 24h. Pseudotyped virus particles were titered on 293T-ACE2 cells by limiting dilution as previously described (*32*). Pseudotypes were diluted to 500 focus-forming units and then incubated with the sera (1:3 diluted from 1:50) at 37°C. Ultra-LEAF IgG1 isotype (Biolegend) and Bamlanivimab (Lilly) were as negative and positive controls (started with 15ug/ml with 3x dilution). After 1h incubation, 293T-ACE2 cells were inoculated with the virus-antibody mixtures and centrifuged at 1200xG for 1 hour at 4°C. Cells were transferred back to the incubator and luciferase was measured at 24 hours post-infection (Bright-Glo^™^ Luciferase Assay System, Promega). Values were normalized to those derived from wells with pseudovirus but without sera (100% infection). Data were input and analyzed by Prism 9. AUC of percentage of infection from each vaccinated and infected were counted and the difference of significance were analyzed by one-way ANOVA test.

## AUTHOR CONTRIBUTIONS STATEMENT

M.L. conceived and designed the study. S.N.S., S.B., M.L. S.F. and B.G performed experiments, analyzed and collected data. M.L. assembled the figures. E.N, K.Z, and J.R. collected and provided serum from patients.

## ACKNOWLEDGMENTS

This work was supported by WSU and the Paul G. Allen School for Global Health (ML BG, SNS, SF, SB). Funding for the clinical study specimens was provided under NIH Project U54CA260581-01 (JM, KZ, EN).

## ADDITIONAL INFORMATION

The authors declare no competing interests.

